## Supplemental Methods for "Translational efficiency across healthy and tumor tissues is proliferation-related"

| REAGENT | SOURCE | IDENTIFIER |
|---|---|---|
| Antarctic phosphatase | New England BioLabs | Cat#M0289 |
| T4 Polynucleotide Kinase | New England BioLabs | Cat#M0201 |
| ProtoScript II Reverse Transcriptase | New England BioLabs | Cat#M0368 |
| miRNeasy Mini kit | Qiagen | Cat#217004 |
| 15% TBE-Urea Gels | NOBEX, Invitrogen | Cat#EC6885BOX |
| RNeasy MinElute Cleanup Kit | Qiagen | Cat#74204 |
| QIAquick PCR Purification Kit | Qiagen | Cat#28106 |
| **CELL LINES** | **SOURCE** | **IDENTIFIER** |
| BJ/hTERT | Gift from Anders H. Lund laboratory (Disa Tehler). | N/A |
| HeLa | ATCC | CCL-2 |
| HEK293 | ATCC | CRL-1573 |
| HCT116 | ATCC | CCL-247 |
| MDA-MB-231 | ATCC | HTB-26 |
| **SOFTWARE** | **SOURCE** | **IDENTIFIER** |
| BBMap [v38.22] | Bushnell B. | https://sourceforge.net/projects/bbmap |
| FastQC [v0.11.4] | Andrews S. | https://www.bioinformatics.babraham.ac.uk/projects/fastqc |
| SAMtools [v1.3.1] | (51) | http://samtools.sourceforge.net |
| tRNAscan-SE [v2.0] | (47) | http://lowelab.ucsc.edu/tRNAscan-SE |
| BEDtools [v2.27.1] | (55) | https://bedtools.readthedocs.io/en/latest |
| Segemehl [v0.3.1] | (48) | https://www.bioinf.uni-leipzig.de/Software/segemehl |
| Picard [v2.18.17] | Broad Institute | https://github.com/broadinstitute/picard |
| GATK [v3.8] | (49) | https://software.broadinstitute.org/gatk |
| GSEA [v3.0] | (50) | https://http:// |

| | | |
|---|---|---|
| | | software.broadinstitute.org/gsea |
