## Supplemental Figures for "Translational efficiency across healthy and tumor tissues is proliferation-related"

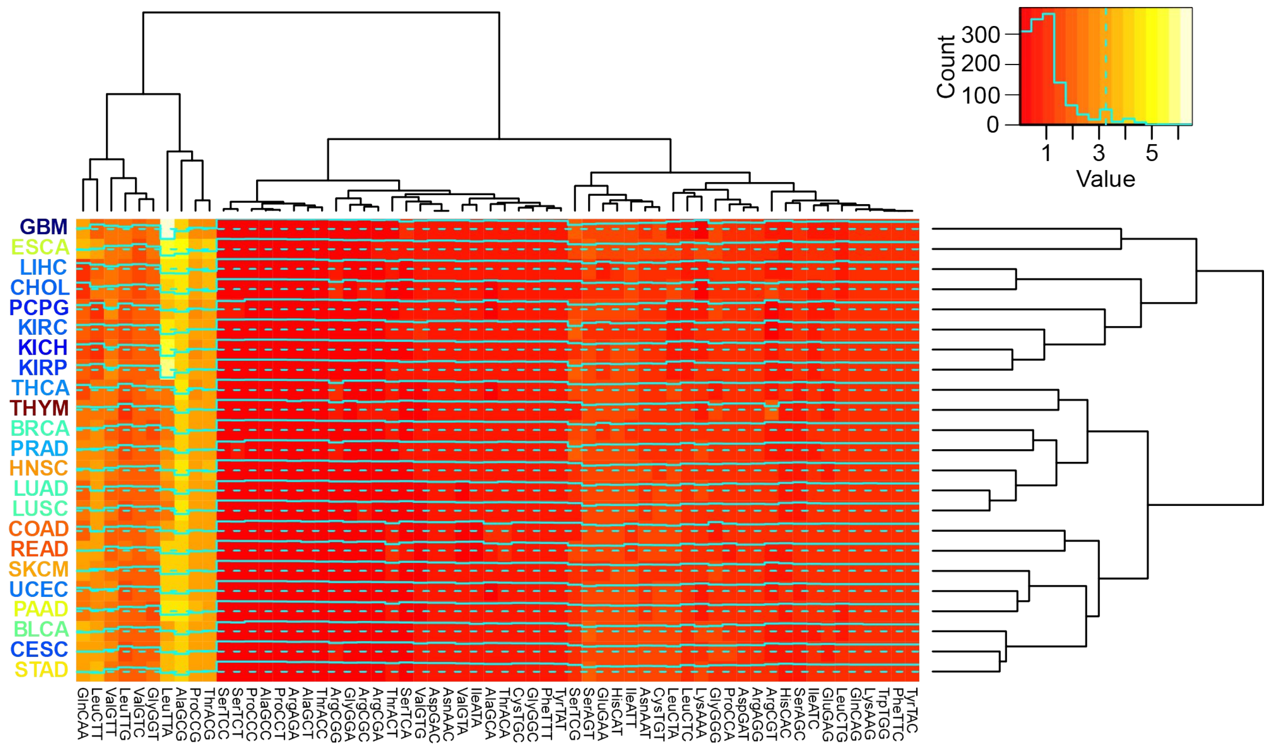


**Figure S1. Medians of Relative Translation Efficiencies weights (RTEw) across all TCGA tissues, related to Figure 3.** The dendrograms show a hierarchical clustering among tissues (right) and among RTEw (top). The tissue labels correspond to the average Ki67 expression. Refer to Supplementary Table 3 for full cancer type names.

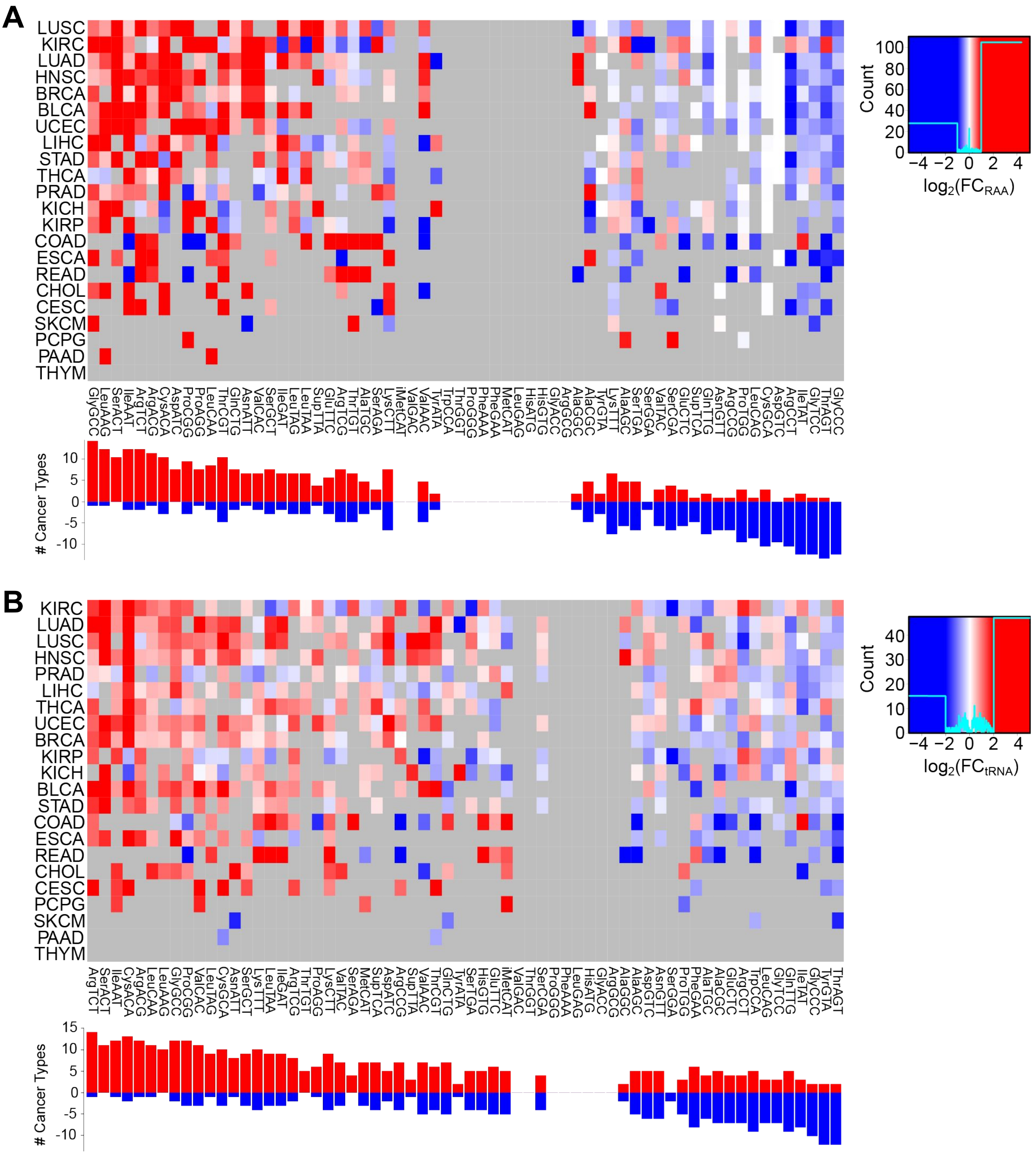


**Figure S2. Differential expression of tRNAs between healthy and tumor samples across 22 cancer types, related to Figure 4.** (A) tRNA expression is measured as Relative Anticodon Abundance and the fold change is $\log_2(RAA_{Tumor}/RAA_{Healthy})$. (B) Absolute tRNA differential expression by $\log_2(tRNA_{Tumor}/tRNA_{Healthy})$. Only significant differences are colored, which are determined using a two-tailed Wilcoxon rank-sum test and corrected for multiple testing by FDR. Refer to Supplementary Table 3 for full cancer type names.

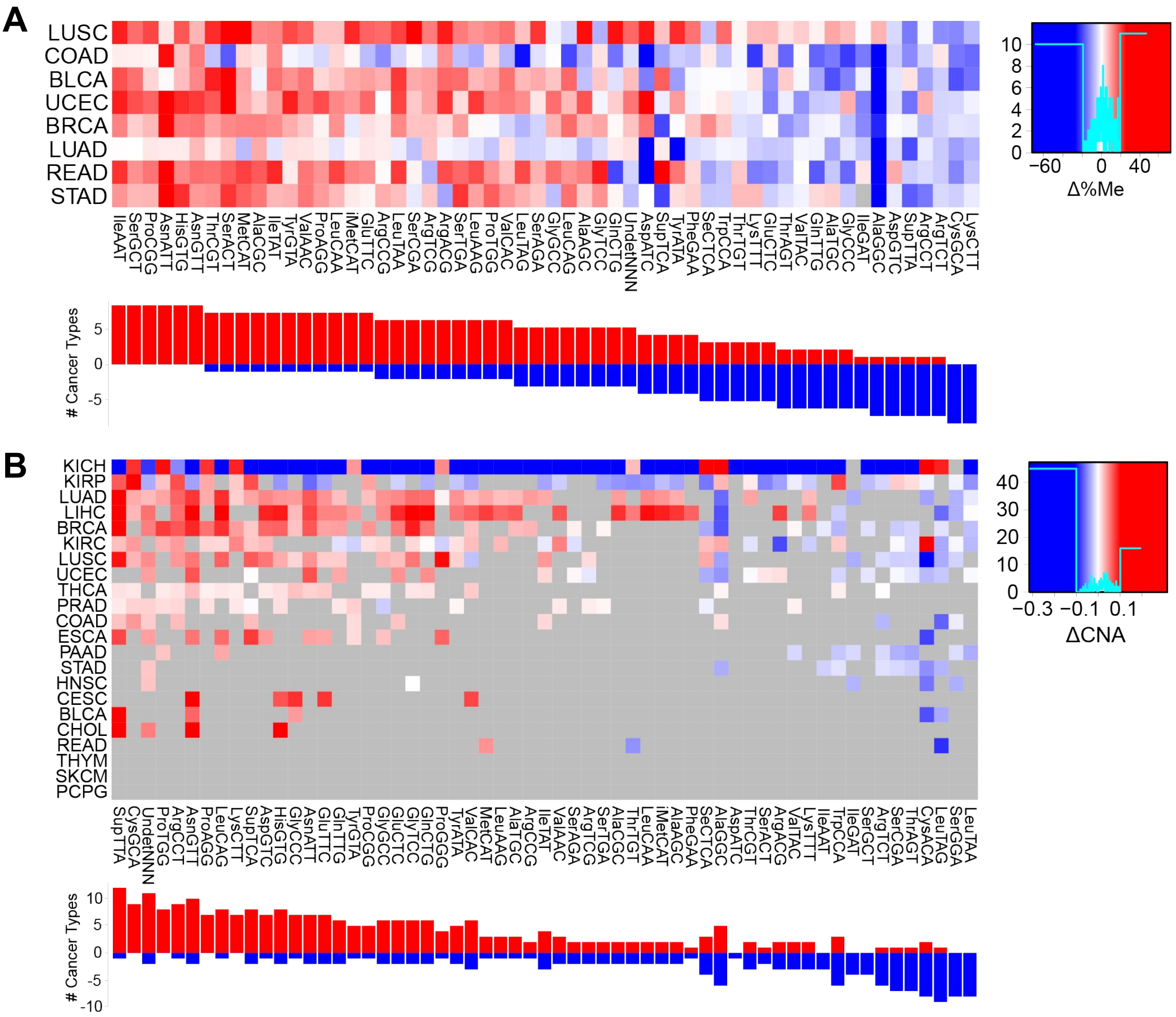


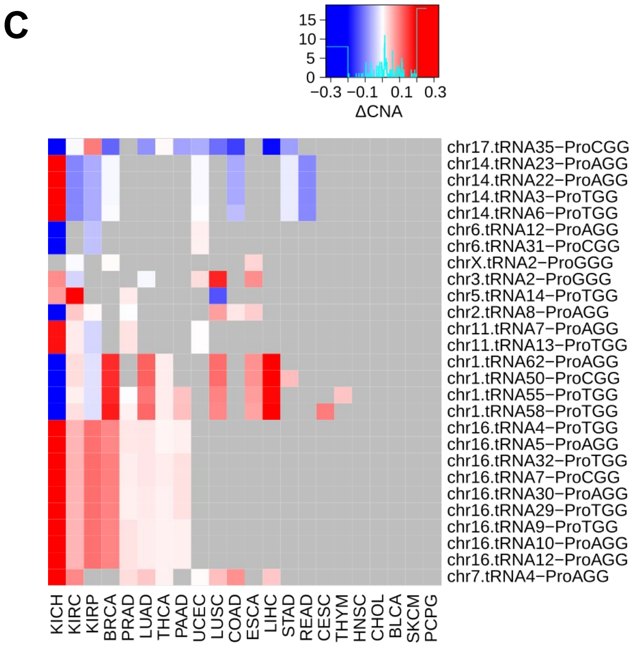


**Figure S3. Differential methylation and copy number between healthy and tumor samples of tRNA genes, related to Figure 5.** (A) Differential average methylation (bisulfite sequencing) for the isoacceptors, averaged over all genes that have a certain anticodon. The difference is measured by $\Delta\%Me=(\%Me_{Tumor}-\%Me_{Healthy})$. (B) Differential average gene copy number for the isoacceptors, averaged over all genes that have a certain anticodon. The difference is measured by $\Delta CNA=(CNA_{Tumor}-CNA_{Healthy})$. Only significant differences are colored, which are determined using a two-tailed Wilcoxon rank-sum test and corrected for multiple testing by FDR. (C) Differential gene copy number between healthy and tumor samples of genes expressing proline tRNAs, as measured by ΔCNA. Only significant differences are colored, which are determined using a two-tailed Wilcoxon rank-sum test and corrected for multiple testing by FDR. Refer to Supplementary Table 3 for full cancer type names.
